# COVIDrugNet: a network-based web tool to investigate the drugs currently in clinical trial to contrast COVID-19

**DOI:** 10.1101/2021.03.05.433897

**Authors:** Luca Menestrina, Chiara Cabrelle, Maurizio Recanatini

## Abstract

The COVID-19 pandemic poses a huge problem of public health that requires the implementation of all available means to contrast it, and drugs are one of them. In this context, we observed an unmet need of depicting the continuously evolving scenario of the ongoing drug clinical trials through an easy-to-use, freely accessible online tool. Starting from this consideration, we developed COVIDrugNet (http://compmedchem.unibo.it/covidrugnet), a web application that allows users to capture a holistic view and keep up to date on how the clinical drug research is responding to the SARS-CoV-2 infection.

Here, we describe the web app and show through some examples how one can explore the whole landscape of medicines in clinical trial for the treatment of COVID-19 and try to probe the consistency of the current approaches with the available biological and pharmacological evidence. We conclude that careful analyses of the COVID-19 drug-target system based on COVIDrugNet can help to understand the biological implications of the proposed drug options, and eventually improve the search for more effective therapies.

## Introduction

The outbreak of the COVID-19 pandemic caused by SARS-CoV-2 at the beginning of 2020 has shocked the population worldwide. One year later, (February 2021) about 110 million confirmed cases of COVID-19 have been reported by WHO included more than 2.4 million deaths (https://covid19.who.int/). As expected, in such a mankind threatening situation, the scientific community put in place a great effort to help countering the spread of the virus, as evidenced among the other things by the huge number of papers dealing with various aspects of the disease appeared in the literature. For instance, the LitCovid literature hub^1^ has collected more than 100,000 articles (February 18, 2021 access) covering arguments categorized as overview, disease mechanism, transmission dynamics, treatment, case report and epidemic forecasting.

As regards the COVID-19 treatment, the race to the vaccine against SARS-CoV-2 started immediately after the isolation of the viral genome^2^ and gave the first results as soon as December 2020. Moreover, despite the exploration of different approaches like, e.g., the infusion of plasma from human survivors^3^, the pharmacological option, namely small molecule drugs and antibodies, is being actively pursued. However, the route to a new drug is long and costly, and the classical drug discovery pipeline is not compatible with the needs of rapid intervention on a population of millions of patients. At the moment, a viable alternative seems to be the repurposing of known drugs^4^, i.e., the use for the treatment of COVID-19 of drugs currently on the market for different therapeutic purposes.

Known drugs that are currently in clinical or pre-clinical study for the treatment of COVID-19 are aimed either at inhibiting viral or human targets involved in some of the processes of viral entry and replication, or at treating inflammation and tissue injury consequent to the viral infection^5,6^. Even though it might seem that a direct antiviral approach could lead to a straightforward solution, only few of the existing antivirals have performed well in the clinic so far. On the other hand, a number of drugs used for the most disparate therapeutic indications and entered into clinical trials even with an uncertain rationale^7^ are showing preliminary promising results. However, as it has been observed^8^, a real “repurposing tsunami” has invested the biomedical community, so much so that today it is difficult not only to keep track of the results of the trials, but also to follow the new proposals.

With the aim of helping researchers navigate the sea of outcomes and reports coming from the studies on COVID-19, some institutions and companies have developed online platforms that collect and organize both literature and data, eventually providing free access to the latter. For example, the already mentioned LitCovid hub^1^ (https://www.ncbi.nlm.nih.gov/research/coronavirus/) is a daily updated source of relevant articles retrieved from PubMed. Other platforms dealing with data on drugs and chemicals, like, e.g., CHEMBL^9^ (https://www.ebi.ac.uk/chembl/), PubChem^10^ (https://pubchem.ncbi.nlm.nih.gov/), or DrugBank^11^ (https://www.drugbank.ca/), have introduced special sections dedicated to COVID-19-related information. In addition, more specialized resources have appeared on the web to help accessing and analyzing COVID-19 data, mainly in the fields of epidemiology, genomics, interactomics, and, to a lesser extent, pharmacology. In this class of web tools, it is worth mentioning CORDITE (CORona Drug InTEractions database)^12^, a web interface that provides a database of potential drugs, targets, interactions, and relative publications obtained from a manually curated selection of literature sources. With the same purpose of facilitating the data analysis, the COVID-19 Drug and Gene Set Library was built as an online collection of COVID-19 related drugs and genes^13^. A comprehensive critical review on this kind of web tools has recently been published by Mercatelli et al.^14^.

Considering the great amount of valuable scientific information that has already been produced and published, and that will be presumably produced for some time more on COVID-19 related topics, it could be useful to look at the whole scenario of results, to foster the acquisition of that knowledge that can only emerge from consideration of both the totality and the complexity of data. In other words, and limiting ourselves to the pharmacological treatment issue, one might think of presenting and analyzing the information on proposed drugs in a way that takes into account not only the different types of data (chemical, biological, genomic, etc.), but also the relationships among them, that is on a network basis. The context is that of network medicine^15^. An attempt in this direction has recently been proposed by Korn et al.^16^, who developed a knowledgebase and an online platform (COVID-KOP) to integrate the existing biomedical information with the newly acquired knowledge on COVID-19. By means of this web tool, one can easily produce an aggregate graph connecting, e.g., COVID-19 phenotypic features to a drug studied for treating the disease, through the genes known to be linked to both. Still in the context of network medicine, CoVex is another platform that offers the user the possibility to explore the SARS-CoV-2 virus-host-drug interactome for drug repurposing aims^17^. In addition, we want to mention CovMulNet19^18^ that at present looks like the most thorough network-based tool allowing to integrate the available genotypic and phenotypic information on COVID-19, like, SARS-CoV-2 proteins, their human partners, as well as symptoms, diseases, and drugs. Finally, Coronavirus canSAR^19^ is a freely available resource that offers druggable interactomes of SARS-Cov-2 proteins and human proteins, as well as reports about 3D structures, drugs, and clinical trials.

In a specifically drug-focused context, the network medicine approach assumes the overcoming of the old “one drug, one target, one disease” concept in favor of a more outright “multi-drug, multi-target, multi-disease” approach^20^. The exploration of a such complex system of interactions can be aided by the construction of a drug-target network^21^. In reference to the COVID-19 case, drug-target networks have been previously built and examined^22–24^ especially with the aim of repurposing already approved drugs.

Here, we present the COVID-19 Drugs Networker (COVIDrugNet: http://compmedchem.unibo.it/covidrugnet), a web application that offers a different point of view on anti-COVID-19 drugs by allowing a network-based analysis of the DrugBank dataset of potential repurposed drugs currently in clinical trial. The freely accessible application automatically retrieves the data from DrugBank, builds the drug-target network, and allows the user to carry out some basic network analysis. Moreover, we show how, using COVIDrugNet, some peculiar aspects of the proposed pharmacological options against COVID-19, in terms of substances, targets, and their interrelationships can be revealed. Although what is reported here is an instant analysis based on current data, the continuous updating of COVIDrugNet will allow us to follow the future development of the drugs proposed for the treatment of the disease, thus providing an always updated view of the COVID-19 system pharmacology.

## Results and Discussion

### COVID-19 Drugs Networker

The COVID-19 Drug Networker (COVIDrugNet, Figure 1) is a web tool designed for the exploration of the landscape of the drugs currently in clinical trials to combat the SARS-CoV-2 infection. The web app is based on a network approach that supports both visualization and analysis of the complex scenario of repurposed drugs for the COVID-19 and related conditions. The core of the web tool are the interactive networks and the additional features that allow one to explore drug and target data, as well as network properties. The main network is a bipartite Drug-Target network (DT, Figure 2a), in which the nodes are drugs and targets, and are connected if a relation between them is reported in DrugBank. Since bipartite networks are usually investigated by compressing their information into two monopartite networks called projections^25^, COVIDrugNet provides two of such networks only having drugs or targets as nodes: in the following, we refer to them as Drug and Target projections, (DP and TP, respectively; Figures 2b and 2c).

**Figure 1.**
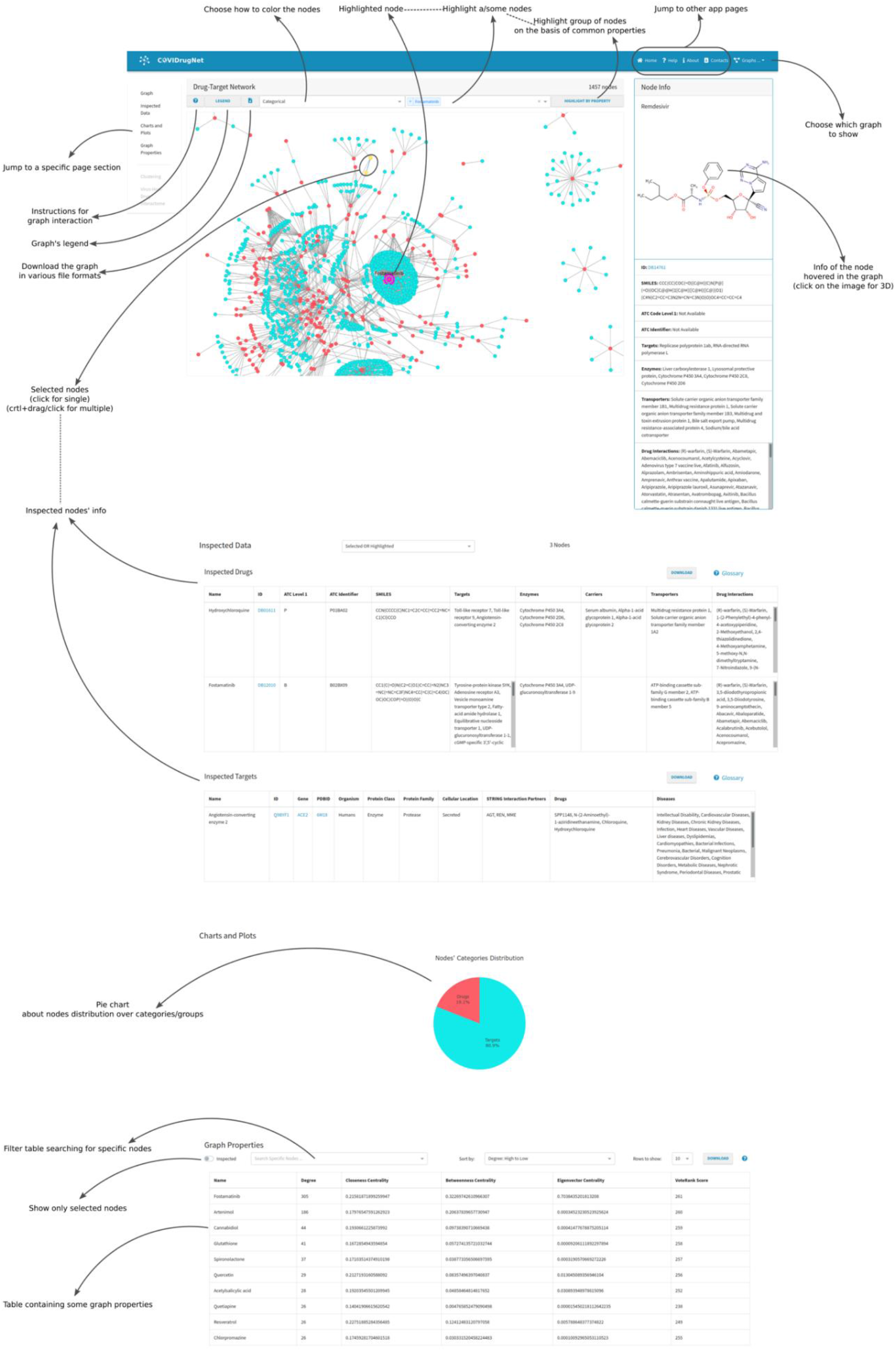
The COVIDrugNet web tool. A screenshot of the main block of the Drug-Target Network page. It displays the fundamental features accessible in the web tool that allow the user to inspect the network and its properties.

**Figure 2.**
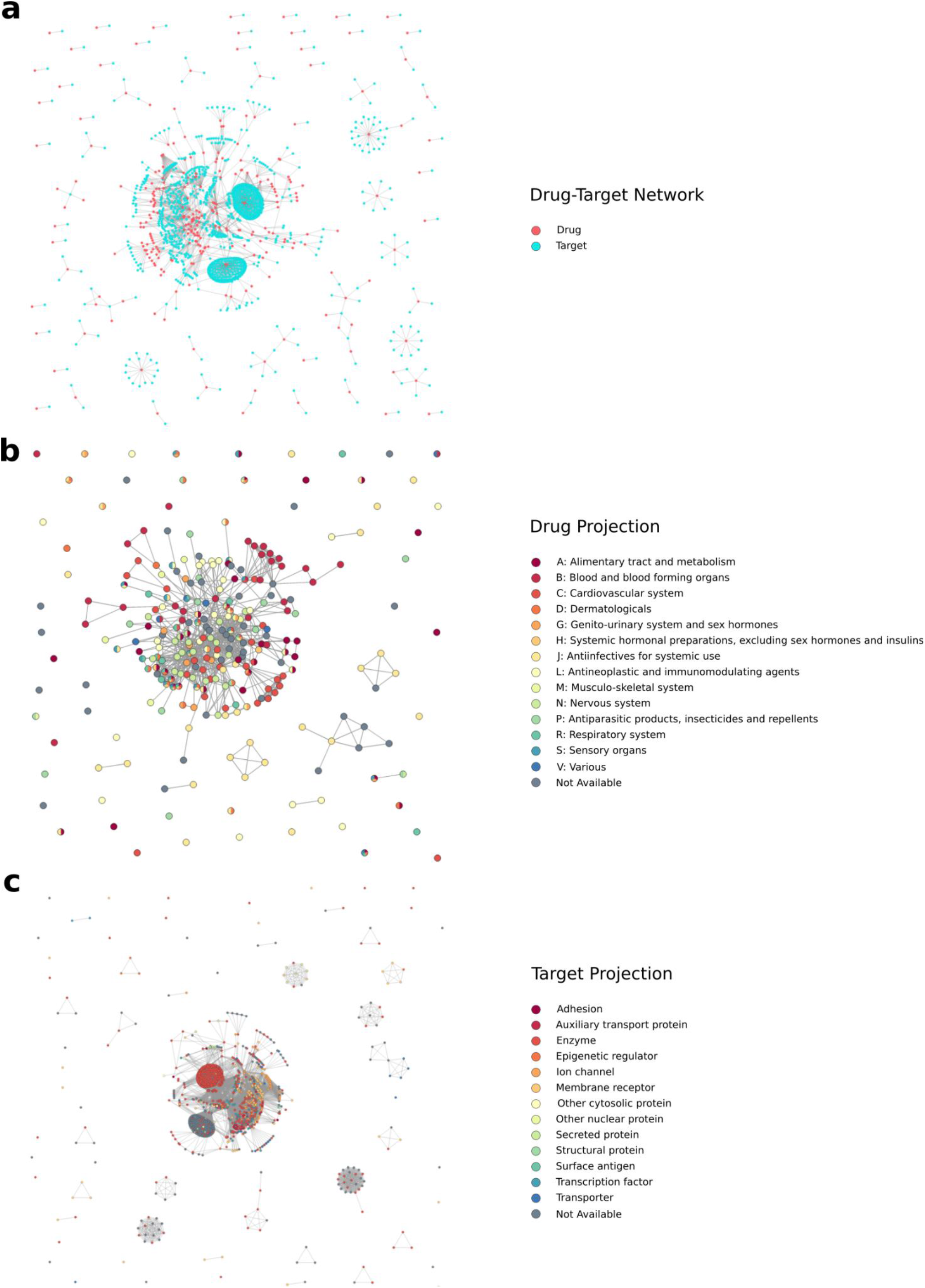
COVIDrugNet Networks. The three networks generated and available for inspection in COVIDrugNet. (a) Drug-Target Bipartite Network. It is the main network, and it is built connecting drugs currently in clinical trial present in the COVID-19 Dashboard of DrugBank^11^ and their reported targets. The red nodes are drugs, and the light blue ones are targets. (b) Drug Projection. It is built from the Drug-Target network and contains only drugs. The nodes are color coded on the basis of their first level ATC codes (retrieved from Drugbank^11^). (c) Target Projection. It is built from the Drug-Target network and contains only targets. The nodes are color coded according to their protein class (retrieved from ChEMBL^9^). The networks were generated by means of the Python package NetworkX^49^.

As regards the user interface, it is basically divided into the main and the *Advanced Tools* blocks. The first one allows users to immediately access the main body of information, capturing the holistic view of the current drug repurposing status for COVID-19. However, a more in-depth examination of the data is possible, by taking advantage of some more specialized graph analysis tools provided in *Advanced Tools*.

In detail, the main block includes the graph, and the *Charts and Plots* and *Graph Properties* sections (Figure 1). As mentioned before, the heart of each web page is the interactive graph with its related information box (*Node Info*) that provides a summary documentation of single drug/target nodes hovered over or individually selected. The box contains links to some databases providing the available information related to individual properties of both drugs and targets. In addition, a multiple node selection brings up the *Inspected Data* hidden section that displays detailed information of the selected nodes in a tabular format. By the way, networks and tables can be downloaded in different formats to allow an external analysis of the data.

Node coloring options are provided, useful to visualize some node attributes related to therapeutic, biological, or network-based features. For instance, the user can decide to color the nodes according to the Anatomical Therapeutic Chemical (ATC) code in the DP, or protein family, protein class or cellular location in the TP. Moreover, in the drug-target network and in both projections, it is possible to color the nodes based on some network attributes - i.e., degree, centrality measures or node grouping - considering the entire graph or the major component. To examine all these properties at a glance, the web tool also provides the *Chart and Plot* section, in which the pie charts - or bar chart in the case of the ATC code coloring option - are updated accordingly to the node coloring option to show the relative proportions between the values of that property. In this area of the projection web pages, the web tool also provides the plot of the nodes’ degree distribution. Among the graph interactive features, the *Highlight a node* dropdown menu is useful to find nodes by name, and the button *HIGHLIGHT BY PROPERTY* allows a customized filtering on node properties to highlight and/or download a specific node selection. In the *Graph Properties* section, some centrality measures useful to analyze the network topology are displayed in a downloadable table. A short explanation of each computed property is provided in a Glossary in the Help page.

Regarding the *Advanced Tools* block, it contains three sections: *Clustering*, *Advanced Degree Distribution* and *Current Virus-Host-Drug Interactome*. The *Clustering* section is dedicated to the node grouping analysis carried out through different methods (see Nodes Grouping section in Methods). In particular, we thought it could be of interest to examine the grouping of the nodes in the projection graphs, as, e.g., in perspective it might reveal possible trends in the selection of drugs to be repurposed or privileged areas of intervention in the biology of the infected cells. To this aim, the web app allows for three different techniques of investigation of the networks partitioning: spectral analysis combined with K-means clustering^26^, Girvan-Newman^27^ and greedy modularity community detection^28^ methods. The plot in this *Clustering* section reports either the eigenvalues distribution used in the application of the spectral clustering method, or the modularity trend in the Girvan-Newman community detection method. Both plots are interactive and allow the user to choose the level (number) of grouping.

The *Advanced Degree Distribution* section presents an interactive chart of the degree distribution and some of its possible distribution fittings compared to those of an Erdős-Rényi equivalent graph (see Degree Distribution Fitting section in Methods).

Finally, the *Current Virus-Host-Drug Interactome* section displays a bipartite network built on the basis of experimental studies and checked for protein targets present in the DT network (see below for details).

### Graphs Analysis

In Figure 2, the graphs representing the networks generated by COVIDrugNet are displayed. The DT network is a disconnected network with a large connected component accounting for 83.2% of nodes (1212 out of 1457). This structure reminds that of the general drug-target network reported elsewhere^21^, where most drugs have more than one target and several drugs can share the same target(s). However, from inspection of the graph, it immediately appears that there are two drug nodes that heavily affect the network topology by showing an exceedingly high degree compared to all other nodes: Fostamatinib and Artenimol, having 305 and 186 direct neighbors, respectively. For both drugs, this reflects a number of reported targets that is considerably higher than the average (<7), being 6.9 and 4.2 times higher, respectively, than that of Cannabidiol that, with 44 targets, is the third in rank for the highest number of neighbors in the DT network. Indeed, these two drugs show a peculiar behavior strongly affecting the network structure not only in the DT, but consequently also in the TP graph where they cause the formation of two highly intra-connected clumps of nodes. To take this aspect under consideration and possibly clarify its role in respect to the topology of both the whole drug-target network and the projections, in the following, we compared the results of the network analyses carried out on the entire networks and on the graphs containing all nodes except Artenimol, Fostamatinib and their exclusive direct neighbors.

As a first step in the analysis, we tried to assess the character of the monopartite projection networks DP (278 nodes) and TP (1179 nodes), i.e., whether they belong to the random network category or are scale-free. Scale-free networks have a characteristic organization, in which there is a limited number of nodes with a high number of neighbors (called hubs) and an abundance of nodes having a low degree^29^. This arrangement can be found in plenty of real-world networks, from the World Wide Web to citations in science, from social interactions to metabolic maps^29,30^. Both DP and TP show a significant difference from an equivalent (same number of nodes and probability of edge creation) Erdős-Rényi graph^31^ (Figure S1). To further investigate on the scale-freeness of the networks, we considered three properties for each graph: the degree distribution, the relationship between clustering coefficient and degree, and the ability to withstand targeted attacks compared to random failures.

In order to address the scale-free character of both networks by evaluating the fitness of the degree distribution to a power-law, we employed the approach reported by Broido et al.^32^, which applied a previously defined rigorous method^33^. This analysis was carried out on both the entire DP and TP networks and also in cases where Artenimol and Fostamatinib as well as their exclusive direct neighbors were removed.

In the Drug Projections, the degree distribution could be described by a power-law, suggesting that these networks are plausibly scale-free (Figure S2a,b). However, other heavy-tailed distributions cannot be ruled out. The situation for the Target Projections is less clear-cut (Figure S2c,d), at least in the case of the entire network. To advance an explanation for these results, we observe that, these networks are small, such that they would probably not provide enough data for clearly electing a distribution form. Still, they are unequivocally dissimilar to random networks.

The inspection of both the clustering coefficient and the robustness evaluation is best illustrated considering the two projections one at a time.

Looking at the DP network and specifically at its clustering coefficient, it shows a tendency to decrease as the degree increases (Figure S3a,b), implicating the existence of a few hubs connecting peripheral nodes of high degree. Also, there is an evident distinction between the response to a targeted attack and to a random failure^34^ (Figure S4a). In the first case, nodes with the highest degree are progressively removed from the network, causing it to break apart quickly. On the other hand, if the nodes to be dismissed are chosen randomly, the connectedness of the network is almost unaffected. Notably, these findings are strengthened by the fact that carrying out the same investigation on a network from which Artenimol and Fostamatinib are excluded, leads to almost identical results (Figure S4b).

The same examination carried out on the TP network does not yield equally unambiguous conclusions. As stated above, the targets linked to Artenimol and Fostamatinib compose two almost-clique aggregations, which distort the morphology of the network. The relationship between clustering coefficient and degree is strongly dependent on the presence of these two exceptionally connected drugs (Figure S3c,d). When they are not taken into account, the inverse proportionality is fairly visible. Nevertheless, if they are considered, the scatterplot displaying this relationship is warped, due to the formation of two separate but remarkably dense groups representing the targets connected to Artenimol and Fostamatinib. The check of the robustness of the network by comparing the responses to targeted attacks or random failures gives a result that agrees with that obtained from the DP network. The communities related to the two “super-spreaders” simply introduce a delay in the fragmentation of the network, since they are made of a multitude of nodes with equally high degree (Figure S4c,d). Anyhow, this shift does not alter the network robustness to random failures and the susceptibility to targeted attacks.

As a final remark on the networks organization, we stress that all results and conclusions presented here are just a snapshot of the continuously evolving COVID-19 drug repurposing scene, and that it will be worthwhile to follow the time progression of this system. For instance, in the future, the growth of the network could smooth out or even hide the effects of Artenimol and Fostamatinib that now we observe so evidently. In respect of this, we recognize a different response of the DP and TP networks to the influence of these nodes. The former is less affected, since the vast majority of the targets related to both drugs are not shared by others, such that the information related to these proteins vanishes in the projection process. On the contrary, the latter suffers a huge impact, showing a situation that is antithetical to the previous one. Here, the proteins amass together constituting two highly intra-connected jumbles, which are poorly linked to the rest of the network. A continuous growth and the ability of self-organizing are two key features of scale-free networks, which frequently describe real complex systems^29^. These characteristics are shown by both projections, and indeed their scale-freeness is supported by their degree distribution, the relation of clustering coefficient to degree, and their robustness. Mainly due to the influence of Artenimol and Fostamatinib, these properties are manifest in the DP network, but not so neat in the TP one.

### Applications to COVID-19 Repurposed Drugs: Network-based Inferences

To illustrate the capabilities of COVIDrugNet, in the following, we report some example considerations that can be derived from the analysis of the projection graphs of both drugs (DP) and targets (TP).

#### Drugs

Examining the DP network with nodes colored by ATC code (https://www.whocc.no/atc/structure_and_principles/) (Figure 2b) can reveal at a glance which therapeutic areas are mostly covered by the repurposed drugs presently in clinical trials. In the *Charts and Plots* section of the COVIDrugNet Drug Projection page, the nodes categories distribution is shown, from which it appears that all the 14 main anatomical/pharmacological groups (1^st^ level codes) are represented, even though with different numbers of drugs. Not taking into consideration the 49 substances for which an ATC code is not yet reported, the remaining 229 drugs are distributed in three top ranked groups: C (Cardiovascular system), A (Alimentary tract and metabolism), and J (Antiinfectives for systemic use) comprising 40, 38, and 38 active substances, respectively (Figure 3). Then, two other highly populated ATC groups follow: B (Blood and blood forming organs), and L (Antineoplastic and immunomodulating agents) counting 32 and 30 drugs, respectively. By considering the composition of the bars that reports the distribution of drugs in the 3^rd^ level groups for each 1^st^ level ATC code (visible in the web tool), one can have a more detailed picture of the actual pharmacological approaches to COVID-19 treatment. First, it is worth noting that the drugs belonging to the J group are located mostly out of the main connected component of the graph, accordingly to the fact that they share a target with a very small number of other drugs.

**Figure 3.**
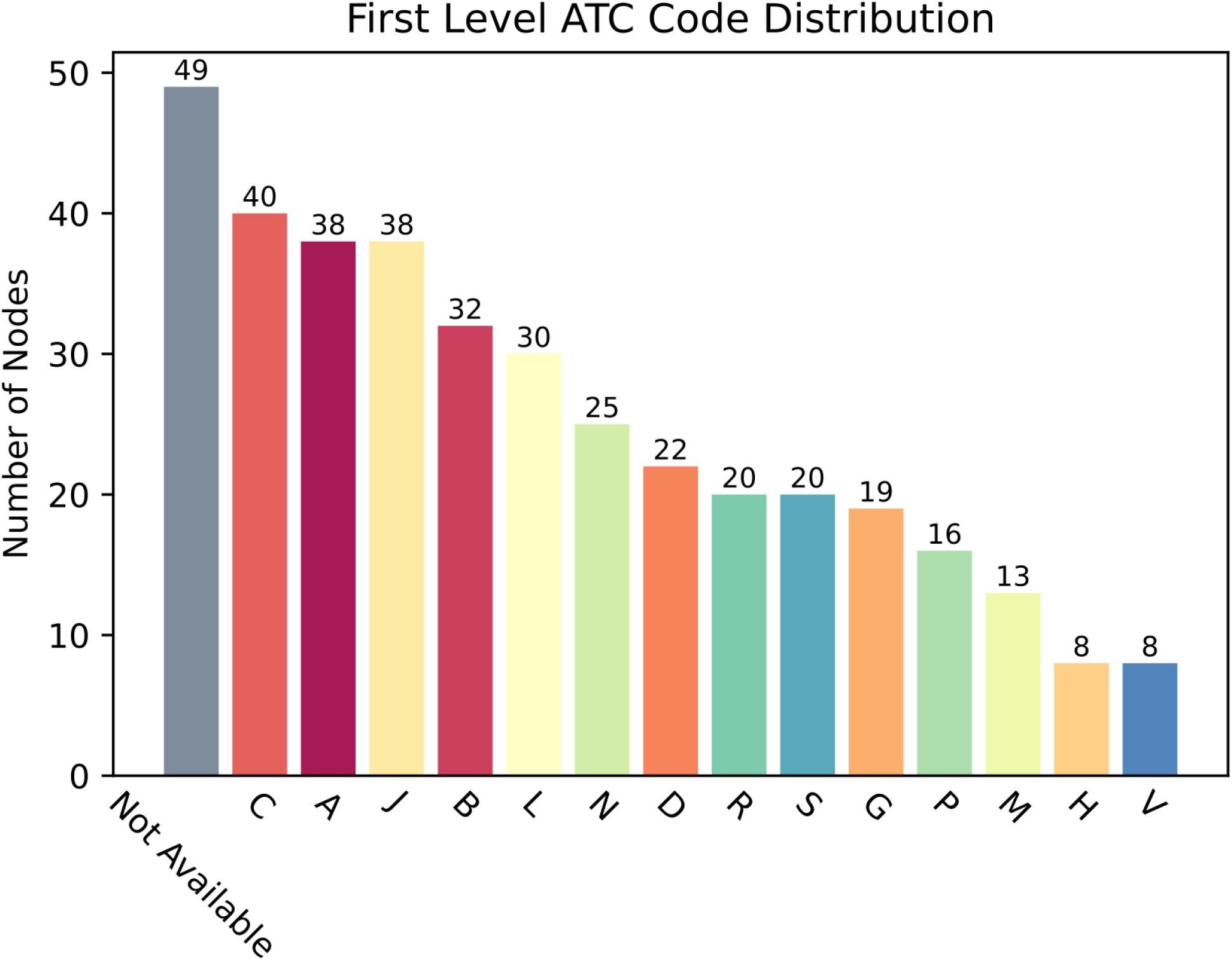
First Level ATC Code Distribution. A bar chart displaying the count of nodes for every first level ATC code (anatomical/pharmacological main group). The total count is higher than the number of nodes in the DP because more than one ATC code can be assigned to a single drug.

Conversely, substances of the A and C groups mostly populate the main connected component, indicating a high level of promiscuity among them as regards the targets. Also, we observe that most drugs classified in the C, A, J, B, L, N, and P groups show just one ATC code, while drugs in D, G, R, S, M, and H belong to more than one ATC group.

Even though the ATC system is not aimed at providing direct therapeutic indications and considering also that more than one code can be assigned to individual medicines, the landscape of pharmacological interventions against the SARS-CoV-2 infection emerging form the DP network appears rather intricate. Overall, it mostly confirms that the drugs in clinical trials are aimed at contrasting both the viral infection process (antivirals in J group, agents acting on the renin-angiotensin system in C group), and its pathological consequences at systemic level (substances in A, B, L, and other groups). These approaches are in line with evidence recognizing that, as the severity of the COVID-19 increases - apparently in consequence of a dysregulated host immune response - various pathophysiological mechanisms are activated leading to hematological (mainly thromboembolic) manifestations and, eventually, multi-organ dysfunctions^35,36^. In addition, bacterial superinfections have been reported in COVID-19 patients, and even though the issue is still debated^37^, antibiotics belonging to the J group are actually in the current treatment guidelines^38^. Indeed, even this brief analysis of the ATC codes distribution among the substances currently in clinical trials highlights a complex and multifaceted drug repurposing scenario consequent to the fact that the COVID-19 is a multi-systemic disease requiring a well-equipped therapeutic armamentarium and possibly a combined poly-pharmacological intervention^39^. However, as it appears from watching the DT network of Figure 2a, some of the drugs considered for therapy are reported to hit such a great number of targets, that it becomes difficult to identify a precise mechanistic hypothesis at the basis of their use. On the other hand, it is conceivable that the promiscuity of these drugs might help counteract the multiple pathological manifestations consequent to the SARS-CoV-2 infection, as well as the viral infection process itself. For example, in the case of Fostamatinib, in the next section, we show that it is able to hit multiple targets involved in critical viral processes, a mechanism that might be at the basis of its favorable pre-clinical profile^40^.

#### Targets

The TP network of Figure 2c is a targetome that shows the relationships among the targets of the proposed COVID-19 drugs. Here, two nodes (proteins) are linked if they are reported as targets of at least one of the drugs in the DrugBank COVID-19 database. The network is made by 1179 nodes and 70794 edges, and shows a main connected component comprising 1011 nodes (85.8%). Human targets are 1013 (884 in the main connected component).

Looking at this graph provides another point of view on the pharmacological approaches taken to contrast the COVID-19. The network of the targets involved in the action of the drugs in clinical trials helps one to obtain a comprehensive view of the biological processes affected by the action of drugs. Actually, from the analysis of the target proteins and their interactions it could be possible to trace the cellular pathways influenced by drugs. A study in this regard is currently underway.

Instead, starting from the TP network, we carried out a different analysis that took into consideration both the data here presented on repurposed drugs now in clinical trials (a top-down view), and the molecular data on SARS-CoV-2 infection obtained from recent experimental studies and exploited to propose drugs to be repurposed (a bottom-up view). As regards the latter, we refer to the human-virus interactomes developed by Gordon et al.^23^ and more recently by Chen et al.^41^.

These interactomes are protein-protein interaction networks that show which human proteins are bound directly by SARS-CoV-2 proteins to allow the virus to enter into the human cells, replicate, assemble and be released. Both research groups followed an experimental approach to identify the human proteins, using affinity purification (AP), and AP together with proximity labeling-based techniques, respectively, coupled with mass spectrometry. Merging the Gordon and Chen results, we obtained an extended list of 732 human proteins experimentally identified as interactors of the 29 viral proteins. Comparing this list with that of the drug targets of the TP network, we found that only 45 out of the 732 human proteins able to bind the viral ones are present in the TP as reported targets of drugs in clinical trials. In Figure 4, we show the integrated host-virus interactome (also available in the *Advanced Tools* block of COVIDrugNet), where the 45 proteins common to both lists are highlighted (yellow circles). We also checked the DT network of Figure 2a for drugs associated with these 45 targets and found 28 substances acting on them (Table 3). They are shown in the interactome of Figure 4 (green squares) linked to their targets. Note that these 28 substances hit direct neighbors of the viral proteins, thus interfering with the related viral processes. Moreover, we see from Figure 4 that Artenimol and Fostamatinib, seemingly by virtue of their high target promiscuity, are able to hit simultaneously several targets, thus affecting various viral processes and allowing to foresee a better therapeutic efficacy. All in all, if confirmed by clinical results, these would be clear examples of poly-pharmacological multi-target actions exerted by single substances, a nice fit into the paradigm of network pharmacology.

**Figure 4.**
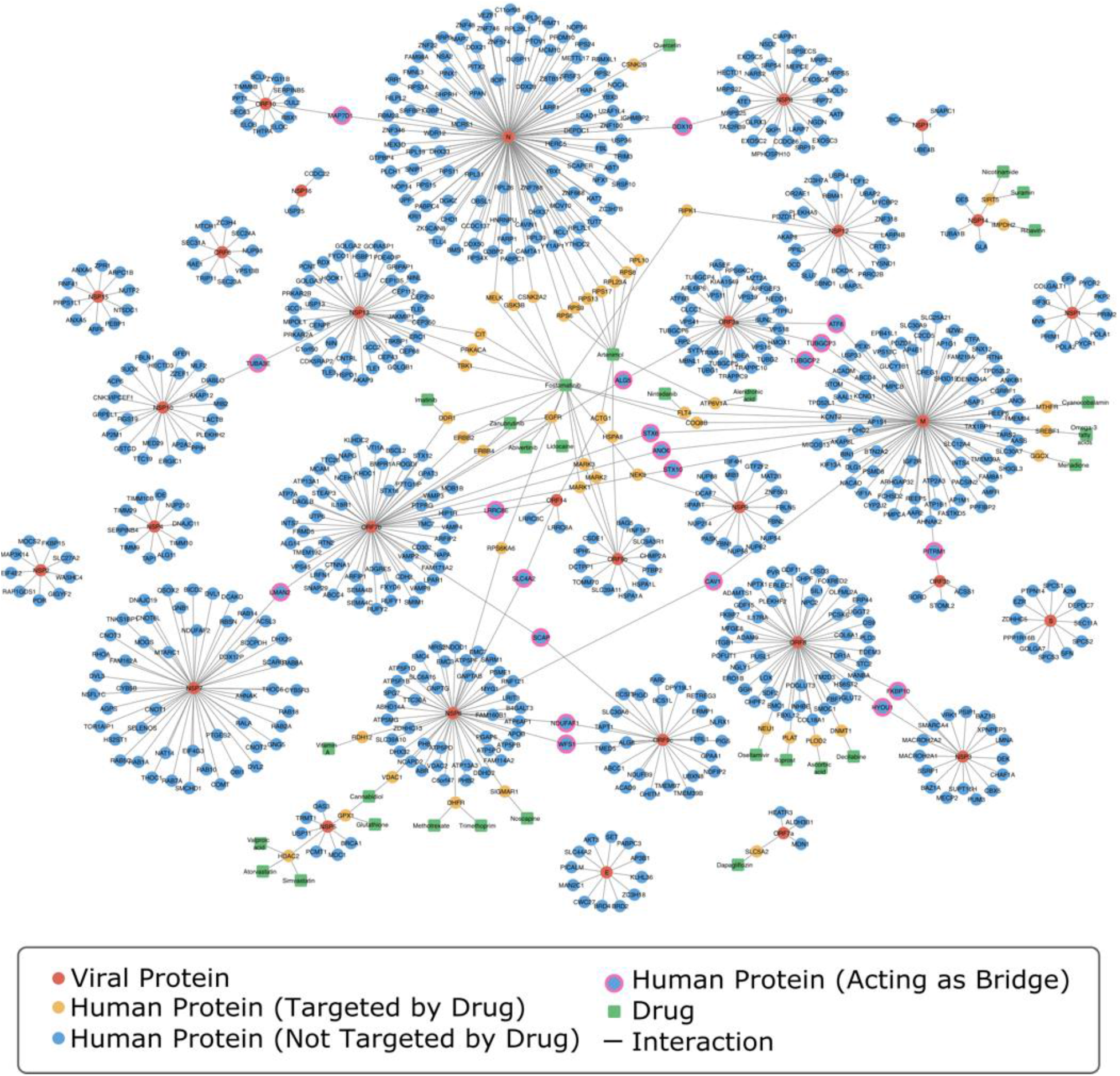
Virus-Host-Drug Interactome. The Virus-Host-Drug Interactome built on the basis of the merged datasets from Gordon et al.^23^ and Chen et al.^41^. Proteins (circles) are displayed in red if viral and in blue if human. The human proteins present in the TP network are shown as yellow circles, and the corresponding drugs currently in clinical trials against COVID-19 as green squares. The human proteins binding more than one viral target are highlighted as blue circles with pink contour. The network visualization was generated through Cytoscape 3.8.2^56^.

Another interesting aspect emerging from inspection of the interactome of Figure 4 is that 20 human proteins (blue circles with pink contour) bind to two viral targets, thus acting as bridges between two node communities and playing a key role in the formation of the large connected component of the graph (Table 4). From a drug discovery perspective, such proteins would be ideal targets to fight the virus, as neutralizing them would help to disrupt the network of interactions necessary to carry on the viral infection and replication processes. Unfortunately, none of these proteins appear in the TP network, implying that there is no substance targeting them among those listed in the DrugBank database of repurposed drugs presently in clinical trial. However, we browsed some databases (DrugBank^11^, DrugCentral^42^ and ChEMBL^9^) in the search for bioactive substances reported to bind these 20 proteins and found that 4 of them are reported as targets of known drugs (Table 5). As can be seen from Table 5, many of the drugs listed therein have not yet been considered for therapy, while some of them are already in clinical trial for COVID-19 treatment even though their action on the proteins in the interactome is not reported in DrugBank. The former ones could be further possible candidates for COVID-19 drug repurposing in light of their ability to interfere with more than one process critical for the virus.

## Limitations

Our study is not exempt from some drawbacks that are common in data analysis, and regard mainly the data availability and quality. We based COVIDrugNet on the DrugBank Dashboard dedicated to COVID-19 pandemic, and although this public and free resource is known for the high reliability of the datasets, missing data or delayed updating can occur. This is evident for some drugs under clinical trial shown in Table 5 that have known targets yet not reported in their DrugBank file. Moreover, not all the drugs or proteins investigated here are completely characterized and classified, and this adds some uncertainty and noise to our results. Also, some bias could be incorporated in the knowledge we started from. For instance, the number of targets associated to a specific drug could considerably depend on the amount of research carried out on that medicine rather than on the actual biological interactions it has. This issue could be partially mitigated by a more extensive integration of data from a wider variety of databases. Very similar considerations can be drawn on the other databases that we exploited to retrieve auxiliary data: STRING^43^, DisGeNet^44^, SWISS-MODEL^45^, RCSB-PDB^46^, UniProt^47^ and ChEMBL^9^. Furthermore, as mentioned in the “Methods” section, the drug-target network was built considering only protein targets, hence nucleic acid targets were not included. However, biomolecular targets other than proteins are a minority^48^ and this led us to not integrate them.

Despite the massive efforts of the scientific community, SARS-CoV-2 and COVID-19 continue to be largely puzzling. Experimental assays are the solid ground on which we all start to build our hypotheses, yet also these investigations may have bias and a moderate amount of uncertainty. We have to keep this into consideration, when examining the merged interactomes by Gordon et al.^23^ and Chen et al.^41^ given their considerable difference. Additionally, the identification of a protein-protein interaction in vitro unfortunately does not guarantee that the same interaction occurs also in vivo.

## Conclusions

The COVID-19 pandemic poses a huge problem of public health that requires the implementation of all available approaches to contrast it, and drugs are one of them. In this context, we observed an unmet need of depicting the continuous evolving scenario of the ongoing drug clinical trials through an easy-to-use freely accessible online tool. Starting from this consideration, we developed COVIDrugNet, a web app that allows one to watch and keep up to date on how the drug research is responding with its arsenal of known repurposed drugs to the health threat represented by the SARS-CoV-2 infection. We have shown some examples of how one could explore the whole landscape of medicines currently in clinical trial and try to probe the consistency of actual treatments with the biological evidence being accumulated on the virus infection and its systemic pathological consequences in humans. The complex network of protein targets affected by the repurposed drugs can be confronted with the host-virus interactome, and this may offer new hints on drugs currently in use or to be proposed for clinical investigation. From this comparison, we have been able to single out some human proteins that contact two viral counterparts, and that might be possible new targets for anti-COVID-19 drugs. Finally, given that, as already noticed by others^7^, several treatments proposed for COVID-19 are still lacking a known mechanism of viral inhibition or even a pharmacological rationale, careful analyses of the drug-target data as those reported in the present work might help to understand the molecular implications of these pharmacological options, and eventually improve the search for more effective therapies.

## Methods

### Data Acquisition

The set of drugs in clinical trial for the treatment of COVID-19 (659 at January 14^th^ 2021) was retrieved from the dedicated web page of DrugBank (https://www.drugbank.ca/covid-19). Both experimental unapproved substances, and drugs in clinical trials were considered, and duplicates were removed (more than one trial is going on for some drugs). The set was also filtered for both the number of heavy atoms (to exclude inorganic compounds), and the availability of data (a drug was not added if it was not present in the PubChem database). This cleaning step reduced the number of drugs considered to 375. From the same site and from PubChem, we gathered some features related to structure, as well as pharmacological classification, pharmacodynamics, and pharmacokinetics of each drug (Table 1). Drugs for which no targets were reported in DrugBank were discarded (278 remaining). As regards the drug targets, they were retrieved from DrugBank, and in this case we collected some information on classification, biology, and pharmacology of each protein. A detailed description of the features and the data sources are reported in Table 2.

**Table 1.** Drugs Features.

| Feature | Description | Source |
| --- | --- | --- |
| ID | DrugBank unique identification code | DrugBank<sup>11</sup> |
| SMILES | The chemical structure string notation for drugs. SMILES were recovered from PubChem if available, otherwise from Drugbank | Pubchem<sup>10</sup><br>DrugBank<sup>11</sup> |
| ATC code level 1 | The broad-based level of the ATC classification system identifying the fourteen anatomical/pharmacological groups | DrugBank<sup>11</sup> |
| ATC identifier | ATC code | Drugbank<sup>11</sup> |
| Targets | Entities to which the drug binds or interacts with, resulting in an alteration of their normal function and thus in desirable therapeutic effects or unwanted adverse effects | Drugbank<sup>11</sup> |
| Enzymes | Proteins that facilitate a metabolic reaction that transforms the drug into one or more metabolites | Drugbank<sup>11</sup> |
| Carriers | Proteins that bind to the drug and modify its pharmacokinetics, e.g., facilitating its transport in the blood stream or across cell membranes | Drugbank<sup>11</sup> |
| Transporters | Proteins that move the drug across the cell membrane | Drugbank<sup>11</sup> |
| Drug Interactions | Drugs that are known to interact, interfere or cause adverse reactions when taken with this drug | Drugbank<sup>11</sup> |

**Table 2.** Targets Features.

| **Feature** | **Description** | **Source** |
| --- | --- | --- |
| Gene | Short identifier of the unique gene name | DrugBank<sup>11</sup> |
| Organism | Organism where the protein comes from | DrugBank<sup>11</sup> |
| Cellular Location | The protein cellular location | DrugBank<sup>11</sup> |
| Drugs | List of known drugs related with the protein (e.g., agonists, antagonists, inhibitors...) | DrugBank<sup>11</sup> |
| ID | UniProt unique identification code | DrugBank<sup>11</sup> |
| STRING Interaction Partners | Known and predicted protein-protein interactions (both physical and functional) only in Homo Sapiens and with a minimum score of 0.95 | STRING<sup>43</sup> |
| Diseases | Disease groups with an Evidence Index of 1 (see https://www.disgenet.org/dbinfo#section44 for more information) | DisGeNET<sup>44</sup> |
| PDBID | Protein Data Bank identification code (the structure with the best resolution) | SWISS-MODEL<sup>45</sup> |
| Protein Classification | The first and the second level of Protein Target Classification are named Protein Class and Protein Family respectively | ChEMBL<sup>9</sup> |

**Table 3.**
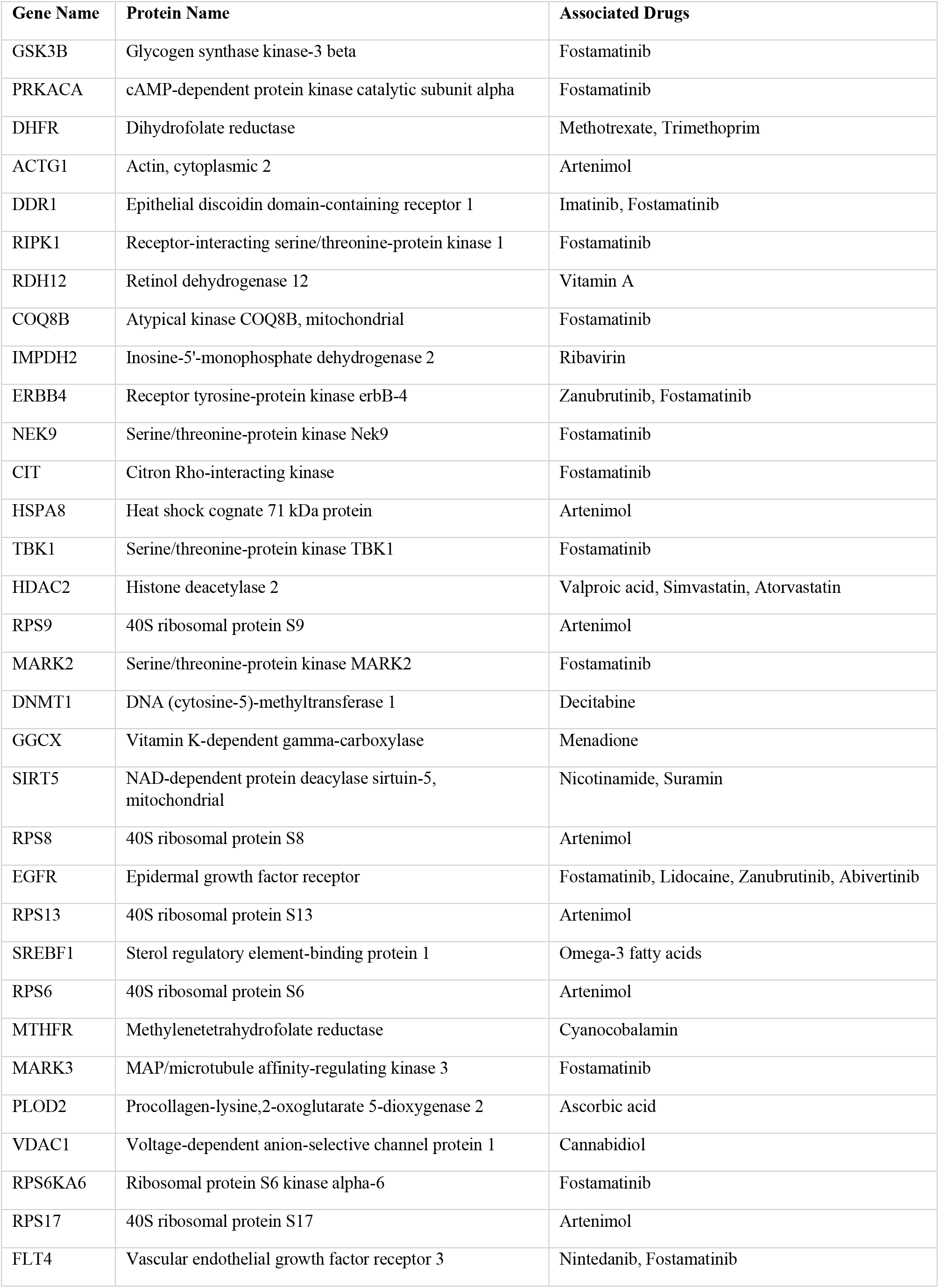
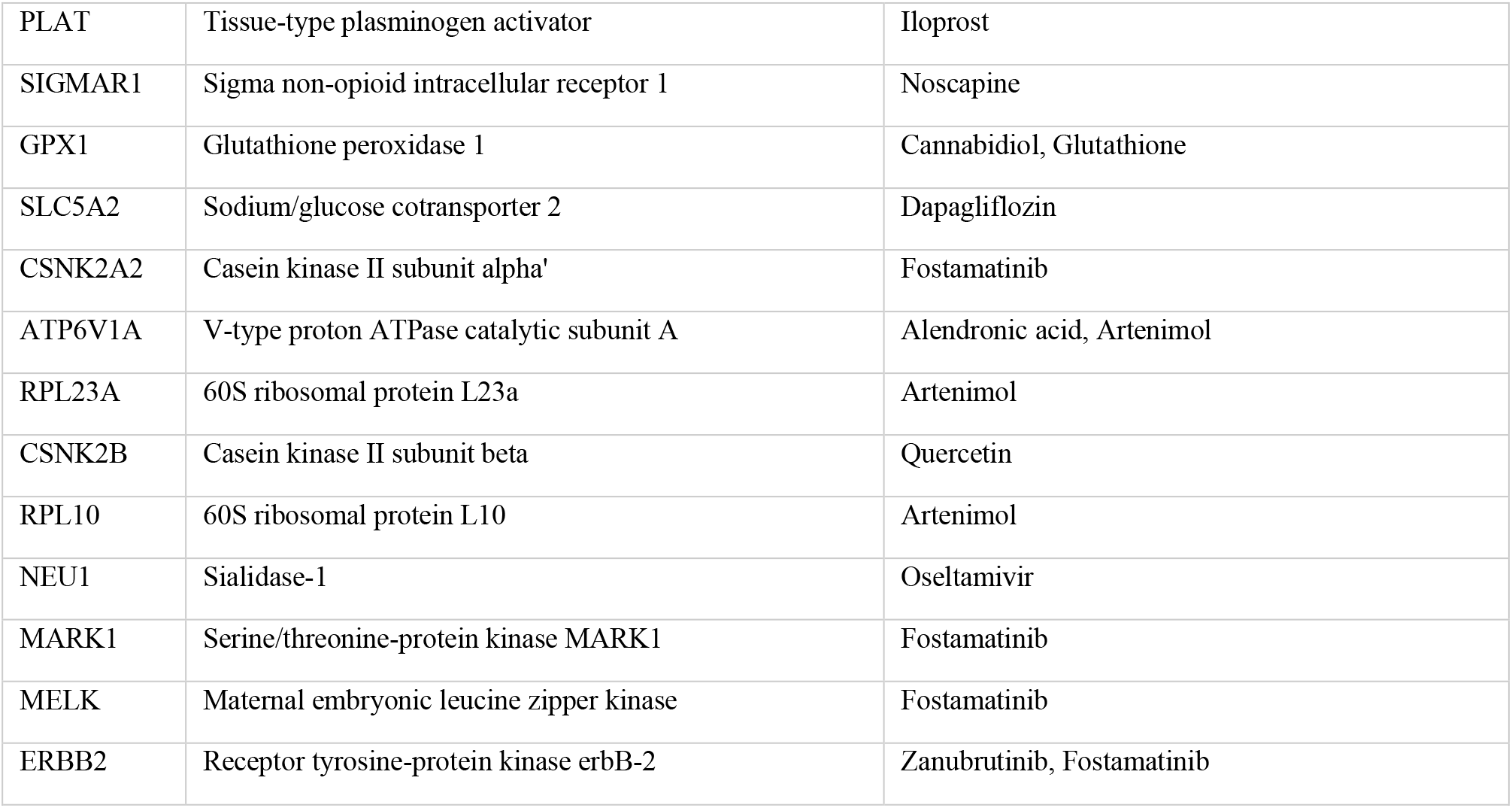
Protein-Drug Associations for Common Targets between the Virus-Host Interactome and the Drug-Target Network.

| Gene Name | Protein Name | Associated Drugs |
| --- | --- | --- |
| GSK3B | Glycogen synthase kinase-3 beta | Fostamatinib |
| PRKACA | cAMP-dependent protein kinase catalytic subunit alpha | Fostamatinib |
| DHFR | Dihydrofolate reductase | Methotrexate, Trimethoprim |
| ACTG1 | Actin, cytoplasmic 2 | Arteminol |
| DDR1 | Epithelial discoidin domain-containing receptor 1 | Imatinib, Fostamatinib |
| RIPK1 | Receptor-interacting serine/threonine-protein kinase 1 | Fostamatinib |
| RDH12 | Retinol dehydrogenase 12 | Vitamin A |
| COQ8B | Atypical kinase COQ8B, mitochondrial | Fostamatinib |
| IMPDH2 | Inosine-5'-monophosphate dehydrogenase 2 | Ribavirin |
| ERBB4 | Receptor tyrosine-protein kinase erbB-4 | Zanubrutinib, Fostamatinib |
| NEK9 | Serine/threonine-protein kinase Nek9 | Fostamatinib |
| CIT | Citron Rho-interacting kinase | Fostamatinib |
| HSPA8 | Heat shock cognate 71 kDa protein | Arteminol |
| TBK1 | Serine/threonine-protein kinase TBK1 | Fostamatinib |
| HDAC2 | Histone deacetylase 2 | Valproic acid, Simvastatin, Atorvastatin |
| RPS9 | 40S ribosomal protein S9 | Arteminol |
| MARK2 | Serine/threonine-protein kinase MARK2 | Fostamatinib |
| DNMT1 | DNA (cytosine-5)-methyltransferase 1 | Decitabine |
| GGCX | Vitamin K-dependent gamma-carboxylase | Menadione |
| SIRT5 | NAD-dependent protein deacylase sirtuin-5, mitochondrial | Nicotinamide, Suramin |
| RPS8 | 40S ribosomal protein S8 | Arteminol |
| EGFR | Epidermal growth factor receptor | Fostamatinib, Lidocaine, Zanubrutinib, Abivertinib |
| RPS13 | 40S ribosomal protein S13 | Arteminol |
| SREBF1 | Sterol regulatory element-binding protein 1 | Omega-3 fatty acids |
| RPS6 | 40S ribosomal protein S6 | Arteminol |
| MTHFR | Methylenetetrahydrofolate reductase | Cyanocobalamin |
| MARK3 | MAP/microtubule affinity-regulating kinase 3 | Fostamatinib |
| PLOD2 | Procollagen-lysine,2-oxoglutarate 5-dioxygenase 2 | Ascorbic acid |
| VDAC1 | Voltage-dependent anion-selective channel protein 1 | Cannabidiol |
| RPS6KA6 | Ribosomal protein S6 kinase alpha-6 | Fostamatinib |
| RPS17 | 40S ribosomal protein S17 | Arteminol |
| FLT4 | Vascular endothelial growth factor receptor 3 | Nintedanib, Fostamatinib |
| PLAT | Tissue-type plasminogen activator | Iloprost |
| SIGMAR1 | Sigma non-opioid intracellular receptor 1 | Noscapine |
| GPX1 | Glutathione peroxidase 1 | Cannabidiol, Glutathione |
| SLC5A2 | Sodium/glucose cotransporter 2 | Dapagliflozin |
| CSNK2A2 | Casein kinase II subunit alpha' | Fostamatinib |
| ATP6V1A | V-type proton ATPase catalytic subunit A | Alendronic acid, Arteminol |
| RPL23A | 60S ribosomal protein L23a | Arteminol |
| CSNK2B | Casein kinase II subunit beta | Quercetin |
| RPL10 | 60S ribosomal protein L10 | Arteminol |
| NEU1 | Sialidase-1 | Oseltamivir |
| MARK1 | Serine/threonine-protein kinase MARK1 | Fostamatinib |
| MELK | Maternal embryonic leucine zipper kinase | Fostamatinib |
| ERBB2 | Receptor tyrosine-protein kinase erbB-2 | Zanubrutinib, Fostamatinib |

**Table 4.** Human Proteins that Interact with more than One Viral Protein in the Virus-Host Interactome.

| **Viral Proteins** | **Human Proteins** | **Name** |
| --- | --- | --- |
| NSP13, NSP10 | TUBA3E | Tubulin alpha-3E chain |
| ORF9C, NSP6 | NDUFAF1 | Complex I intermediate-associated protein 30, mitochondrial |
| NSP3, ORF8 | FKBP10 | Peptidyl-prolyl cis-trans isomerase FKBP10 |
| M, ORF3a | ATF6 | Cyclic AMP-dependent transcription factor ATF-6 alpha |
| M, ORF7b | STX10 | Syntaxin-10 |
| ORF7b, ORF14 | LRRC8E | Volume-regulated anion channel subunit LRRC8E |
| M, ORF3a | TUBGCP3 | Gamma-tubulin complex component 3 |
| NSP6, ORF14 | SLC4A2 | Anion exchange protein 2 |
| ORF8, NSP3 | HYOU1 | Hypoxia up-regulated protein 1 |
| M, ORF7b | STX6 | Syntaxin-6 |
| M, ORF3a | TUBGCP2 | Gamma-tubulin complex component 2 |
| ORF10, N | MAP7D1 | MAP7 domain-containing protein 1 |
| N, NSP8 | DDX10 | Probable ATP-dependent RNA helicase DDX10 |
| ORF9c, NSP6 | WFS1 | Wolframin |
| M, ORF5b | PITRM1 | Presequence protease, mitochondrial |
| ORF7b, M | ANO6 | Anoctamin-6 |
| ORF7b, NSP7 | LMAN2 | Vesicular integral-membrane protein VIP36 |
| M, NSP6 | CAV1 | Caveolin-1 |
| ORF9c, ORF7b | SCAP | Sterol regulatory element-binding protein cleavage-activating protein |
| ORF3a, ORF7b | ALG5 | Dolichyl-phosphate beta-glucosyltransferase |

**Table 5.** Known Drugs Targeting Human Proteins that Interact with more than One Viral Protein in the Virus-Host Interactome.

| Human Proteins | Name | Known Drugs |
| --- | --- | --- |
| TUBA3E | Tubulin alpha-3E chain | Podophyllotoxin<sup>a</sup><br>CYT997<sup>b</sup><br>Docetaxel<sup>c</sup><br>Vincristine<sup>c</sup><br>Verubulin<sup>c</sup><br>Indibulin<sup>c</sup><br>Trastuzumab – Emtansine<sup>c</sup><br>Ixabepilone<sup>c</sup><br>Sagopilone<sup>c</sup><br>Eribulin<sup>c</sup><br>Fosbretabulin<sup>c</sup><br>Mirvetuximab – Soravtansine<sup>c</sup><br>Paclitaxel<sup>c</sup><br>Plinabulin<sup>c</sup><br>Polatuzumab – Vedotin<sup>c</sup><br>Vinblastine<sup>c</sup><br>Crolibulin<sup>c</sup><br>Fosbretabulin<sup>c</sup><br>Cabazitaxel<sup>c</sup><br>Davunetide<sup>c</sup><br>Paclitaxel – Poliglumex<sup>c</sup><br>Vinflunine<sup>c</sup><br>Lexibulin<sup>c</sup><br>**Colchicine**<sup>*c</sup><br>Vinorelbine<sup>c</sup> |
| NDUFAF1 | Complex I intermediate-associated protein 30, mitochondrial | **Metformin**<sup>*d</sup><br>NV-128<sup>e</sup><br>ME-344<sup>e</sup> |
| FKBP10 | Peptidyl-prolyl cis-trans isomerase FKBP10 | **Tacrolimus**<sup>*f</sup> |
| ATF6 | Cyclic AMP-dependent transcription factor ATF-6 alpha | Pseudoephedrine<sup>g</sup> |

\*) Drugs currently in clinical trial for COVID-19
a) Retrieved from DrugCentral (https://drugcentral.org/target/Q6PEY2/)
b) Retrieved from DrugBank (https://go.drugbank.com/drugs/DB05147)
c) Retrieved from ChEMBL
([https://www.ebi.ac.uk/chembl/g/#browse/mechanisms of action/filter/target.target ch embl id%3ACHEMBL2095182](https://www.ebi.ac.uk/chembl/g/#browse/mechanisms%20of%20action/filter/target.target%20ch%20embl%20id%3ACHEMBL2095182))
d) Retrieved from DrugCentral (https://drugcentral.org/target/Q9Y375/)
e) Retrieved from ChEMBL
([https://www.ebi.ac.uk/chembl/g/#browse/mechanisms of action/filter/target.target ch embl id%3ACHEMBL2363065](https://www.ebi.ac.uk/chembl/g/#browse/mechanisms%20of%20action/filter/target.target%20ch%20embl%20id%3ACHEMBL2363065))
f) Retrieved from DrugCentral (https://drugcentral.org/target/Q96AY3/)
g) Retrieved from DrugBank (https://go.drugbank.com/drugs/DB00852)

### Networks Construction

We chose to inspect the data in the form of a graph. All networks presented in the web app and in the paper were built by means of the NetworkX software^49^.

### Network Analysis

#### Node Attributes

Some suitable node attributes (Degree, Closeness Centrality, Betweenness Centrality, Eigenvector Centrality, Clustering Coefficient, VoteRank) were calculated through NetworkX. The only property we tweaked was the result of the VoteRank because its algorithm draws up a ranking of nodes based on an iterative voting system^50^ without assigning a specific value to each one of them. Thus, we translated this ranking into a score for each node on the basis of its position and the total number of nodes with the following method:

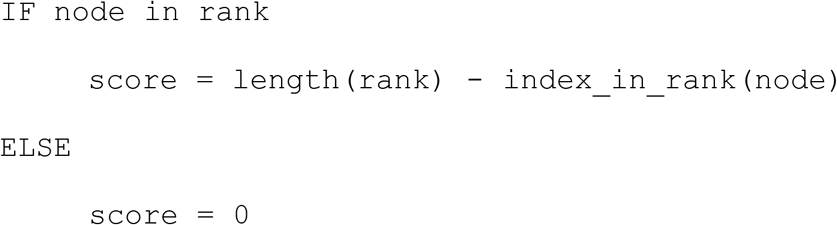

#### Nodes Grouping

Dividing a network into groups, clusters, or communities could be useful to unveil not trivial patterns of interaction. It is accomplished by splitting the network into subgroups that have the fewest possible number of connections between them^51^. In this work, we took advantage of (and provide access to in the webtool) three of the most common algorithms for this purpose: spectral clustering^26^, Girvan-Newman community detection^27^, and greedy modularity community detection^28^.

The first one makes use of the spectrum of the graph Laplacian to convey the information about the graph partition^26^. The division is then carried out on this data by a k-means clustering algorithm (see the Supplementary Information for more detail). In the second case, communities are recognized employing the Girvan-Newman method^27^. It is a hierarchical method based on the progressive removal of the edges with the highest betweenness centrality from the graph, causing it to break into sets of smaller constituents. The partition with the best modularity is shown, but the user can manually choose an arbitrary number of communities in the web tool. The greedy modularity community detection method^28^ pursues the graph division through a bottom-up approach (opposite to the previous one), by exploiting a “greedy” algorithm that progressively associates the nodes into groups that maximize the modularity. It starts with all nodes separated into single communities and recursively merges the couple of them that brings to the highest modularity increasing, until the point that joining two communities would lead to a modularity reduction. These tasks were accomplished through in-house Python scripts, mainly making use of the packages NetworkX and Scikit-learn^52^.

#### Degree Distribution Fitting

A network is commonly considered to be scale-free if the degree distribution of its nodes follows a power-law^29^, which has the form:

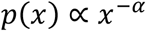

where the scaling exponent α is higher than 1 (usually between 2 and 3) and the degree value x is equal or greater than x_min_ (which is always higher than 1). To the best of our knowledge, the most severe scale-freeness test is presented by Broido et al.^32^ that take advantage of a rigorous mathematical procedure^33^ to assess the validity of a power-law distribution to describe the investigated degrees. Here, we followed their approach probing the fitting of a power-law to the degree distributions of both projected networks DP and TP (with and without the Artenimol and Fostamatinib nodes). As a first step, the parameters of the best fitting power-law are determined (x_min_ with a standard Kolmogorov-Smirnov minimization approach, and then α with a discrete maximum likelihood estimation) employing the Python package Powerlaw^53^. Then, the fitting is evaluated considering the p-value of the Kolmogorov-Smirnov distance (computed with a semi-parametric bootstrap), and of the x_min_ and α (bootstrap). If p ≥ 0.1, the degree distribution is considered plausibly scale-free. Lastly, the chosen power-law distribution is compared to four non-scale-free alternatives (using loglikelihood ratio tests), to evaluate if it is favored over the others. Such alternatives are the exponentially truncated power-law, the exponential, the stretched exponential (Weibull) and the lognormal. This entire procedure was carried out using an in-house Python script, with a large employment of the Python package Powerlaw. A more thorough explanation and method validation are provided in the Supplementary Information.

#### Robustness

Scale-free networks (contrary to random Erdős-Rényi graphs) have an exceptional tolerance against random failures, but at the same time they are very vulnerable to targeted attacks^34^. We investigated the robustness of these networks evaluating their diameter (as a measure of interconnectivity) throughout a process of node removal. We took into account both targeted attacks and random failures and compared the results. In the first case, at every iteration the node with the highest degree was chosen and removed. In the other case, a node was selected randomly and eliminated. In this latter condition, the average of multiple 100 runs was considered in order to avoid misinterpretations induced by a single random choice. This procedure was carried out through an in-house Python script.

### COVIDrugNet Implementation and Deployment

COVIDrugNet is mainly composed by the collector and the web tool itself. Both are written in Python, but the purpose of the former is to collect the data from web databases, build the graphs, compute some properties, and store everything in pickle format. The latter, instead, retrieves the data from the created database and sets up the front-end part of COVIDrugNet with Python Dash^54^.

The web tool deployment was carried out with Apache^55^ through the mod_wsgi interface in an Ubuntu server.

## Supporting information

Supplementary Graph Data

## Acknowledgements

This work was supported by University of Bologna. The authors wish to thank Fabrizio De Ponti for the helpful discussions and Riccardo Ocello for the support in the server management.

## Author contributions

L.M., C.C., M.R. conceived and designed the study. L.M. performed the acquisition, integration of the data, adapted the algorithms for the analyses, carried out the tests, prepared the figures and implemented the web tool, and C.C. performed the web server configuration and set up the interface with the web tool. M.R. was in charge of overall direction, planning, and supervision. All authors provided critical feedback and helped in the interpretation of data, tested and provided original contributions to the improvement of the app, and wrote the paper. All authors read and approved the final manuscript.

## Additional Information

### Data and Code Availability

The full code for the collection, building and analysis of the networks is available in the GitHub repository at https://github.com/LucaMenestrina/COVIDrugNet. It is entirely written in Python and makes use of the packages listed in the Supporting Information. COVIDrugNet is a public web tool available at http://compmedchem.unibo.it/covidrugnet. All data generated or analyzed in this study is publicly available and is included in this article (and its supplementary information files) or on the GitHub repository (in which data will be updated every two weeks). Furthermore, some data is easily downloadable from the web tool itself: all tables in tab-separated values (tsv) format and the networks in various formats (adjacency list, pickle, cytoscape json, graphml, gexf, edges list, multiline adjacency list, tsv, png and jpg).

## Notes

http://compmedchem.unibo.it/covidrugnet/

https://github.com/LucaMenestrina/COVIDrugNet

### Competing Interest Statement

The authors have declared no competing interest.

