## Supplementary Information for "COVIDrugNet: a network-based web tool to investigate the drugs currently in clinical trial to contrast COVID-19"

*

### Degree Distribution Fitting

#### *Fitting the model*

Fitting a power-law distribution means estimating the α parameter for describing the available data as:

$$p(x) \propto x^{-\alpha}$$

The first step to accomplish this goal is to choose the lower limit of the node degrees ($x_{min}$) on which to fit the distribution. We carried out this operation using a Kolmogorov-Smirnov minimization approach[1], which aims to find the fitting with the lowest absolute difference between the cumulative density function of the fitted distribution and the cumulative empirical distribution (both for $x \geq x_{min}$).

The scaling exponent α is instead selected employing the maximum-likelihood estimation technique.[1]

Both computations are executed through the Python package Powerlaw[2].

#### *Evaluating the quality of the fitting*

The obtained fitting is tested taking into consideration the p-value of the Kolmogorov-Smirnov distance (D), of the $x_{min}$ and of the α parameter. The latter two are assessed with a case resampling bootstrap, in which the synthetic samples are built with a random sampling with replacement from the node degrees. On the other hand, the first one (D) is estimated with a semi-parametric bootstrap approach[1]. In this case, the construction of the synthetic sample is done separately (but keeping the same proportion of the empirical data) for the portions of degrees lower and higher than $x_{min}$. The first part is randomly sampled from the fraction of empirical degrees lower than $x_{min}$. The other one, instead, is generated by Powerlaw as a random deviate of the fitted model.

The fitting is considered plausible if the p-value of D is at least 0.1, which means having a 10% probability of false negatives in the case the underlying generative process is indeed scale free.

#### *Comparing the power-law fitting to the ones of other heavy-tailed distributions*

Determining whether the power-law is a plausible distribution is necessary but not sufficient for stating that there are no other models better describing the empirical data. Thus, it is mandatory to compare the fits of different candidate distributions. Here we employed log-likelihood ratio tests[1] for analyzing the power-law, the exponentially truncated power-law, the exponential, the stretched exponential (Weibull) and the lognormal distributions. However, there will always be a distribution that describes the data in a better way (since there are infinite possibilities).

A power-law distribution is part of the so-called heavy-tailed distributions, which, by definition, have the exponential as the absolute minimum alternative[2]. Hence, in order to consider the power-law as a feasible distribution, it has to be at least more likely than the exponential one.

#### *Method implementation validation*

Our implementation of the power-law fitting, its quality evaluation and its comparison to other heavy-tailed distributions were tested on the same datasets on which the Powerlaw package was validated[2]: the frequency of word usage in the novel ‘‘Moby Dick’’ by Herman Melville (for optimal fitting), the number of each neuron connections in *C. elegans* (for moderate fitting) and the number of customers in the United States affected by electricity blackouts (for poor fitting).

**Spectral Clustering**

We first calculated the graph Laplacian[3]

$$L = D - A$$

where D is the diagonal matrix of node degrees and A is the adjacency matrix.

Then, we computed its normalized form:

$$L_{norm} = D^{-1/2} L D^{-1/2}$$

The eigenvalues of the normalized Laplacian matrix and the corresponding eigenvectors convey the intel about the graph partition.[3,4] The connected components are as many as the 0 eigenvalues and are defined by the relative eigenvectors. The remaining eigenvectors designate the subsequent clustering.

We chose the number of clusters taking advantage of the eigengap heuristic.[4] In order to automatize this choice, we devised the following procedure. First, calculate the difference between every consecutive eigenvalue and fit this batch to a half-normal distribution (only the positive part of a normal distribution). Then, compute the percent point function for 99% (the probability of having a difference above this number is 1%) and consider the first occurring eigenvalue with an increase from the previous one higher than the determined threshold (if more consecutive eigenvalues are eligible, the highest one is used). Lastly, choose its index if it is higher than 1 (for the major component) or not equal to the number of components (for the entire graph), else the index of the following occurrence is used. The number of clusters suggested by this method is generally acceptable, but the user is allowed to manually choose it in the webtool. After that, a k-means clustering algorithm on the selected eigenvectors with 100 independent runs (for improving consistency) was used to select the groups.

**Python Modules**

Python modules employed throughout the whole work

- Python 3 (core)
  - pandas (v 1.1.5)
  - numpy (v 1.19.4)
  - os
  - pickle
  - json
  - itertools
  - requests (v 2.25.1)
  - time
  - datetime
  - webbrowser
  - threading
  - matplotlib pyplot (v 3.3.3)
- networkx (v 2.5)
- plotly (v 4.9.0)
- dash (v 1.13.4)
  - dash core components (v 1.10.1)
  - dash html components (v 1.0.3)
  - dash bootstrap components (v 0.10.3)
  - dash cytoscape (v 0.2.0)
  - dash daq (v 0.5.0)

- powerlaw (v 1.4.6)
- PubChemPy (v 1.0.4)
- chembl webresource client (v 0.10.2)
- beautiful soup (v 4.9.3)
- rdkit (v 2009.Q1-1)
- scikit-learn (v 0.14.0)
- tqdm (v 4.54.1)
- visdcc (v 0.0.40)

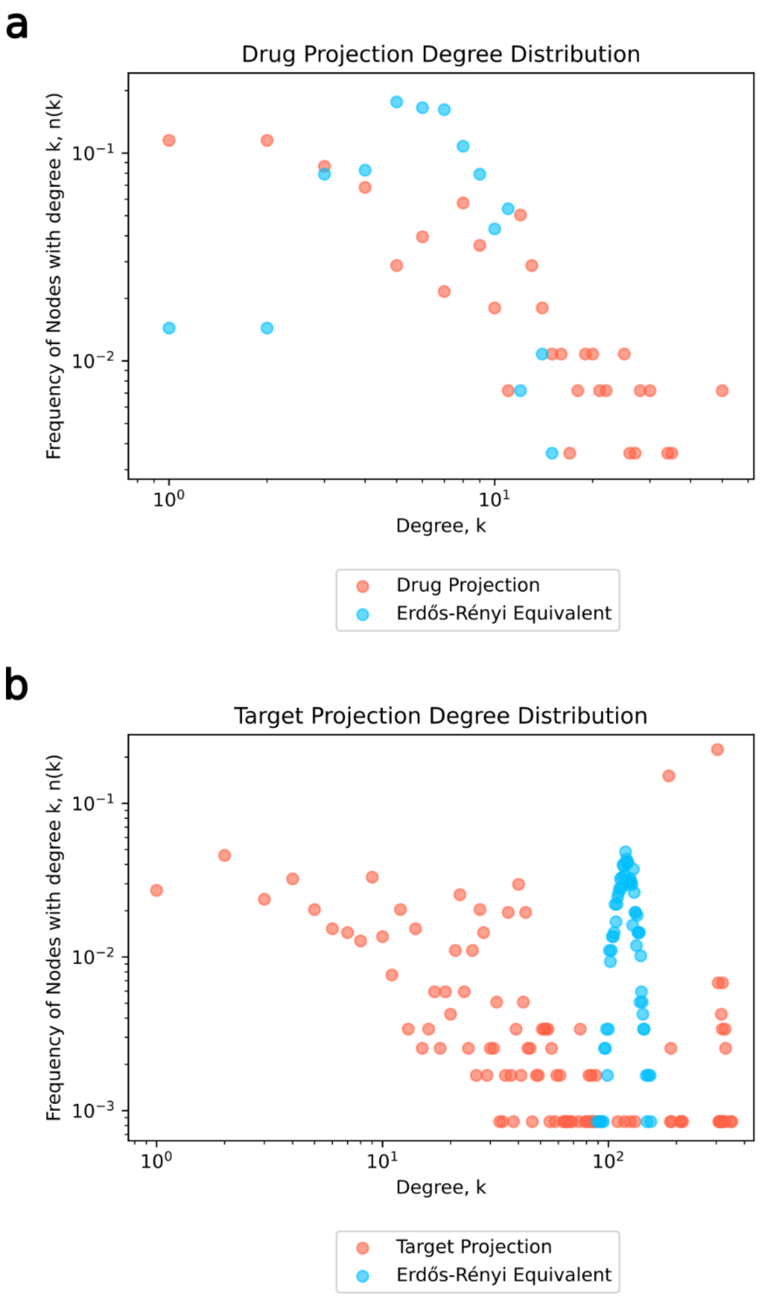


***Figure S1. Degree Distribution***

The degree distributions of both the drug (a) and target (b) projections (red), compared to those of equivalent random graphs (blue). The last ones were generated with the Erdős-Rényi model[5], keeping the same number of nodes and probability of edge creation (ratio between the actual number of edges and the maximum possible edges) of the network they are compared with.

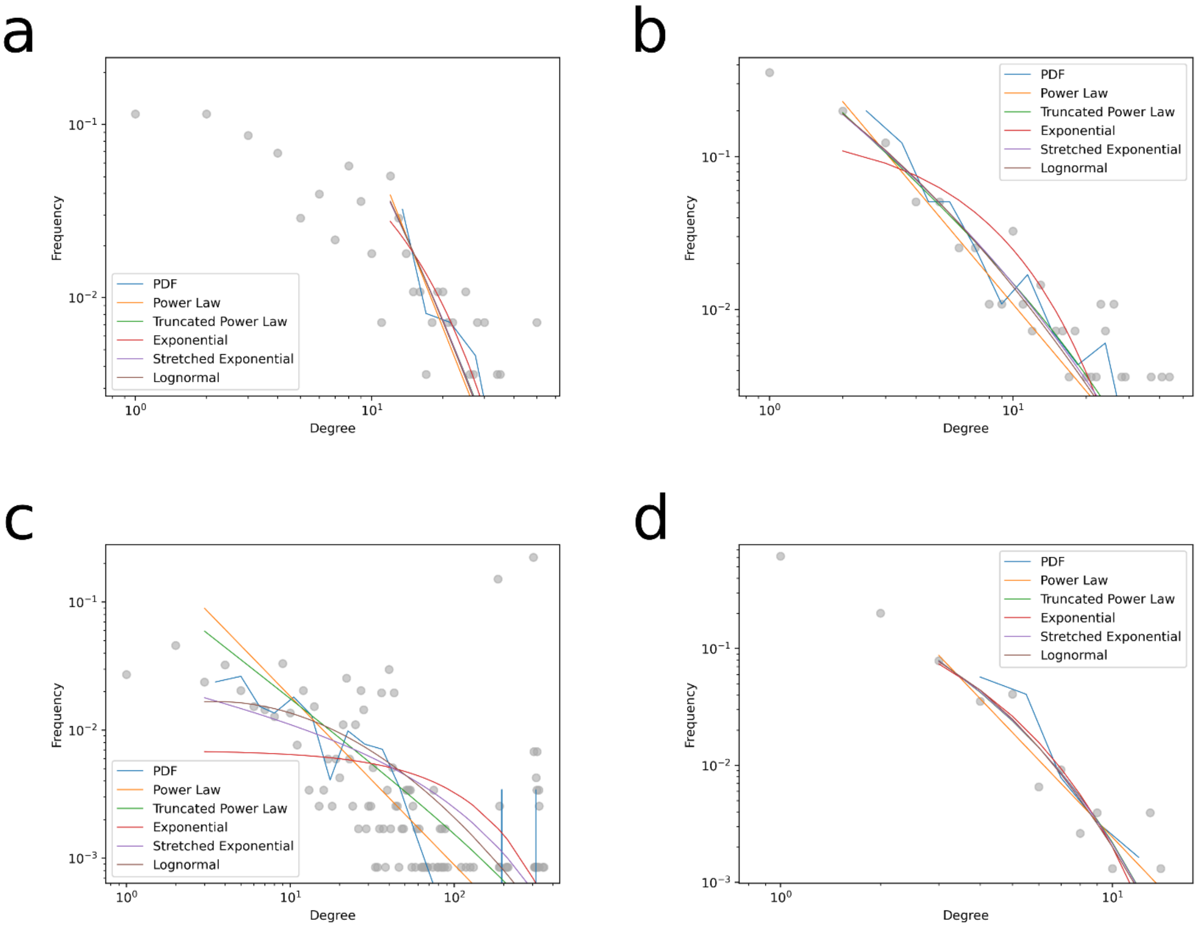


***Figure S2. Degree Distribution Fittings***

The degree distributions of the entire drug projection (a) and entire target projection (c) networks, and of the corresponding graphs in which all nodes except Artenimol, Fostamatinib and their exclusive direct neighbors were present (b and d, respectively). On each distribution the following functions are fitted: power-law (orange), truncated power-law (green), exponential (red), stretched exponential (violet) and lognormal (brown). In every chart the probability density function (PDF, blue) is shown too.

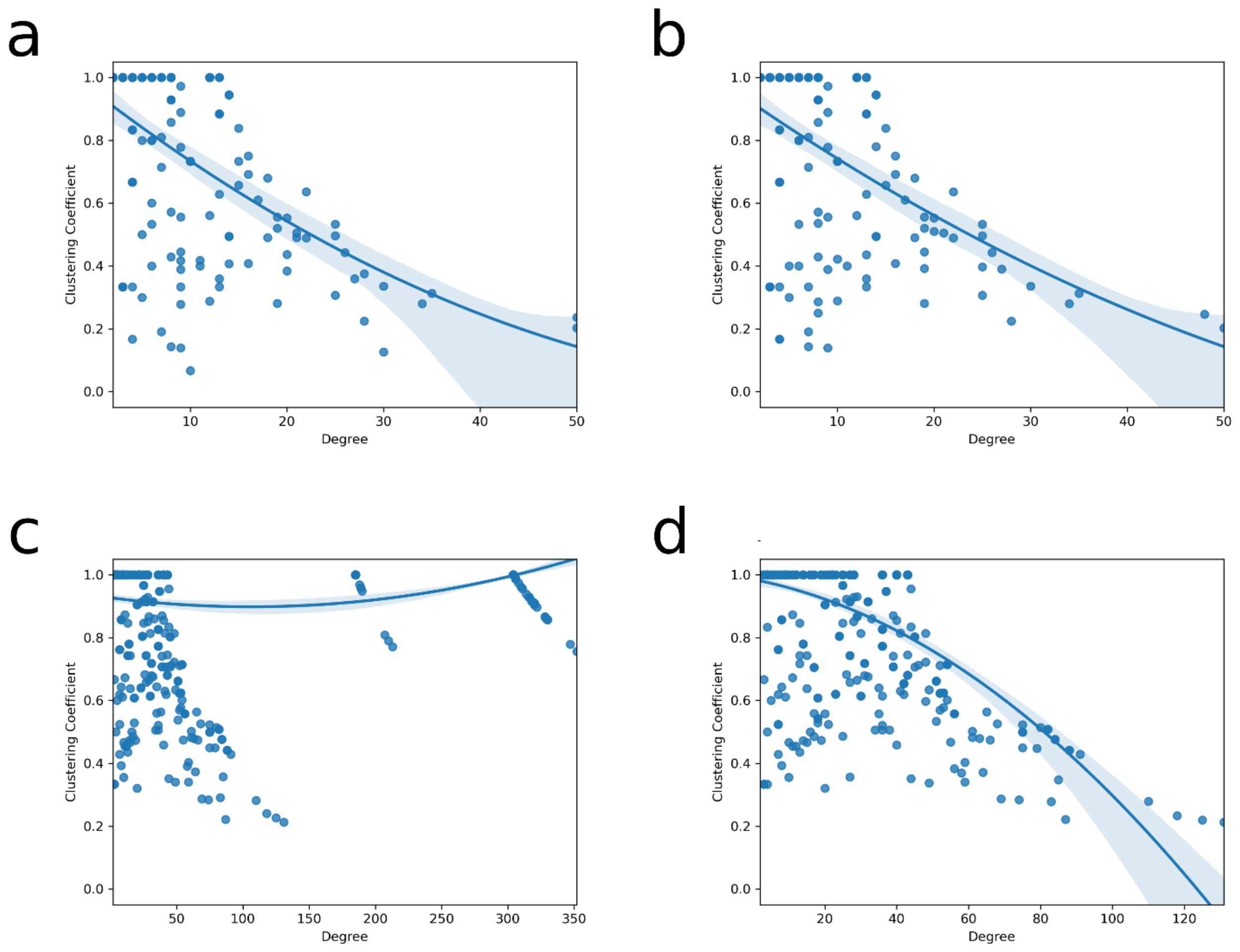


***Figure S3. Clustering Coefficient and Degree Relationship***

The relationship between the clustering coefficient and the degree of nodes in the entire drug projection (a), the entire target projection (c) networks, and the corresponding graphs in which all nodes except Artenimol, Fostamatinib and their exclusive direct neighbors were present (b and d, respectively).

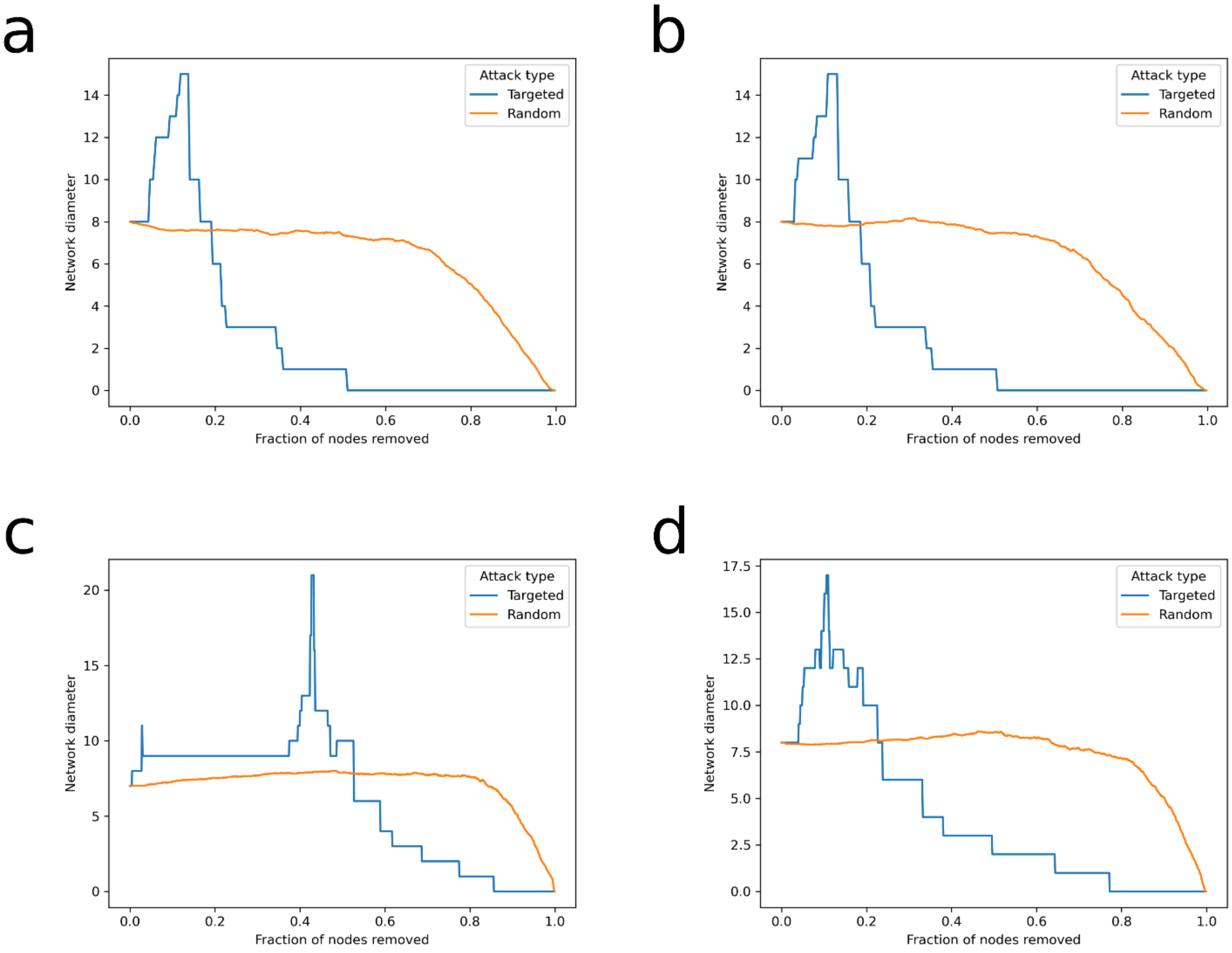


***Figure S4. Network Robustness***

The comparison of the network diameter variation in response to targeted attacks (in blue) and random failures (in orange). The investigation was carried out on the entire drug projection (a), the entire target projection (c) networks, and the corresponding graphs in which all nodes except Artenimol, Fostamatinib and their exclusive direct neighbors were present (b and d, respectively).

| | Drug Projection (Entire) | | Drug Projection | | Target Projection (Entire) | | Target Projection | |
|---|---|---|---|---|---|---|---|---|
| | D | p-value | D | p-value | D | p-value | D | p-value |
| Power-Law | 0.08 | $2.0 \times 10^{-1}$ | 0.05 | $3.1 \times 10^{-1}$ | 0.22 | 0.0 | 0.06 | $5.1 \times 10^{-2}$ |
| | Likelihood-ratio | p-value | Likelihood-ratio | p-value | Likelihood-ratio | p-value | Likelihood-ratio | p-value |
| Truncated Power-Law | -0.49 | $3.2 \times 10^{-1}$ | -8.55 | $3.5 \times 10^{-5}$ | -355.34 | 0.0 | -4.90 | $1.7 \times 10^{-3}$ |
| Exponential | 1.18 | $6.1 \times 10^{-1}$ | 13.60 | $1.1 \times 10^{-1}$ | -365.64 | $2.8 \times 10^{-21}$ | -4.34 | $1.9 \times 10^{-1}$ |
| Stretched Exponential | -0.33 | $6.2 \times 10^{-1}$ | -7.41 | $3.9 \times 10^{-3}$ | -432.81 | $9.6 \times 10^{-63}$ | -4.94 | $3.0 \times 10^{-2}$ |
| Lognormal | -0.26 | $6.3 \times 10^{-1}$ | -6.49 | $6.0 \times 10^{-3}$ | -351.58 | $1.6 \times 10^{-60}$ | -5.03 | $4.2 \times 10^{-2}$ |

***Table S1. Fittings Evaluation***

The table provides the log-likelihood and respective p-value for each function fitted on every analyzed network. For the power-law function, the Kolmogorov-Smirnov distance (D) is provided, and the fitting is considered plausible if the respective p-value is at least 0.1.
